# Adsorption and conformations of lysozyme and *α*-lactalbumin at a water-octane interface

**DOI:** 10.1101/155341

**Authors:** David L. Cheung

## Abstract

As they contain both hydrophobic and hydrophilic amino acids proteins will readily adsorb onto interfaces between water and hydrophobic fluids such as oil. This adsorption normally causes changes in protein structure, which can result in a loss of protein function and irreversible adsorption, leading to the formation of protein interfacial films. While this can be advantageous in some applications (e.g. food technology) in most cases it limits our ability to exploit protein functionality at interfaces. To understand and control protein interfacial adsorption and function it is necessary to understand the microscopic conformation of proteins at liquid interfaces. In this paper molecular dynamics simulations are used to investigate the adsorption and conformation of two similar proteins, lysozyme and *α*-lactalbumin, at a water-octane interface. While they both adsorb onto the interface *α*-lactalbumin does so in a specific orientation, mediated by two amphipathic helices, while lysozyme adsorbs in a non-specific manner. Using replica exchange simulations both proteins are found to possess a number of distinct interfacial conformations, with compact states similar to the solution conformation being most common for both proteins. Decomposing the different contributions to the protein energy at oil-water interfaces, suggests that conformational change for *α*-lactalbumin, unlike lysozyme, is driven by favourable protein-oil interactions. Revealing these differences between the factors that govern conformational change at interfaces in otherwise similar proteins can give insight into the control of protein interfacial adsorption, aggregation, and function.

## I. INTRODUCTION

The adsorption of proteins onto liquid interfaces, such as the air-water or oil-water interfaces, has been harnessed in a number of materials applications. Proteins have long been used as emulsion stabilisers in food technology^1^ and recently a number of novel biosurfactant proteins have been identified that have potential for use in functional foods with attractive properties^2^. Protein layers at fluid interfaces have also been exploited as templates for the growth of inorganic materials through biomineralization^3^. Interfacial adsorption of proteins is also important in a number of biological contexts. Some of the best known examples are biosurfactants^4,5^ which act to reduce surface tension and stabilise foams and emulsions. These include the hydrophobins^6^, amphiphilic proteins that are expressed by filamentous fungi, aiding the formation of aerial structures such as spores or hyphae, ranaspumin, that is used by the tropical frog *Engystomops pustulosus* in the production of foam nests^7^, and latherin^8^ which is found in horse sweat. More sophisticated interfacial function is exhibited by lipases^9^, enzymes that break down fats and which become active only when adsorbed onto oil-water interfaces, and lung surfactant proteins, which fulfil a number of functions including immune response^10^.

When proteins adsorb onto liquid interfaces they often undergo conformational change^11,12^. This is driven by the hydrophobic amino acids, which typically form the core of the protein, partitioning into the hydrophobic medium. For biosurfactants or other proteins that have evolved to function in interfacial environments this conformational change is often an intrinsic part of their function, such as the lid-opening seen in lipases or unhinging transition exhibited by Rsn-2^13^. In the more general case of proteins for which interfacial adsorption is not part of their function this is typically associated with a loss of activity, which limits the applications of proteins in interfacial environments. The unfolding of proteins at interfaces is also responsible for strong, often essentially irreversible adsorption, which can lead to loss of protein material during processing (e.g. in the biopharmaceutical industry).

As understanding the factors that drive interfacial adsorption and conformational change and how to control this process is important in a number of biological and technological areas, this has been the subject of considerable investigation with both experimental and theoretical approaches. X-ray^14^ or neutron reflectivity^15^ can examine the size of proteins, in particular the layer thickness. Neutron reflectivity has investigated the unhinging transition of the biosurfactant Rsn2 at the air-water interface^13^ and the structure of mixed protein layers at interfaces^16^. Information on the orientation of secondary structure elements (helices or strands) at interfaces can be obtained from infra-red spectroscopy. Secondary structure at interfaces has been investigated using refractive index matched emulsion circular dichroism^17^ (RIME CD) and synchrotron radiation circular dichroism^18^ (SRCD). Formation of larger-scale structures has been investigated using AFM, tensiometry, and rheological measurements (e.g. passive particle probe tracking)^19^. While these methods cannot directly investigate protein structure different aggregation behaviour can be related changes in protein conformation.

Complementing these experimental studies a number of groups have used molecular dynamics simulations to investigate protein interfacial adsorption and structure. Many of these have focused on the initial stages of interfacial adsorption, aiming to determine the factors that mediate protein adsorption^20–22^. Molecular dynamics simulations have also been used to investigate the interactions between proteins and inorganic ions at interfaces, giving insight into the process of biomineralization at interfaces^23,24^. Simulations have also been used to determine the adsorption strengths of proteins to liquid interfaces. These have shown that amphiphilic proteins have adsorption free energies ~ 100 *k*_B_*T*^25,26^, essentially showing irreversible adsorption to interfaces. For smaller proteins and peptides this has been measured to be smaller, typically ~ 10 *k*_B_*T*^27,28^, but still significant.

Due to the associated timescales conformational change of proteins at interfaces has been less studied. For small peptides it is possible to examine changes in secondary structure associated with interfacial adsorption within timescales that can be accessed in simulation^29^. To do this for larger proteins it is necessary to use advanced simulation techniques, such as replica exchange^30^ or metadynamics^31^. Recently replica exchange simulations of lysozyme at the water-dichloroethane interface showed that this can adopt a number of conformations of similar free energy^32^. Simulations of peptides derived from myoglobin also found that these exhibit a number of different interfacial conformations, with the preferred orientation for the two peptides investigated being consistent with their differing emulsification behaviour^33^.

To understand protein adsorption and conformations at interfaces it is necessary to have a microscopic picture of this process, for example identifying which regions of the protein undergo changes in structure. While experimental measurements using SRCD or RIME CD can determine the overall amounts of secondary structure in a protein they cannot determine which regions of the protein undergo these changes. Understanding this will give detailed information about the mechanisms of conformational change and how this is related to protein structure. SRCD measurements on a number of proteins have shown that while the secondary structure of proteins is different at interfaces compared to bulk solution, they often retain a large amount of ordered secondary structure^34–36^. In most cases the distribution of this secondary structure and whether this is the same as in solution is not clear. Additionally as previous simulation work suggests that even globular proteins can exhibit a number of distinct conformations at interfaces, experimental measurements are likely to give information averaged over the ensemble of different interfacial conformations.

In this paper the conformations of two proteins, lysozyme (LSZ) and *α*-lactalbumin (*α*La) (Figure 1(a)), at a water-octane interface (which may be considered as an archetypal water-oil interface) are investigated. LSZ and *α*La are homologous proteins; they are of similar sizes (LSZ has 129 residues, *α*La 123) and analysis of their hydropathy plots (Figure 1(b)) show that these have similar patterns of hydrophobic and hydrophilic residues. Previous work using SRCD on these proteins at oil-water interfaces have shown that these exhibit differing changes in secondary structure at interfaces, with the proportion of *α*-helix increasing for *α*-lactalbumin^35^ and decreasing for lysozyme^36^. Molecular dynamics simulations are used to investigate how these adsorb at interfaces, and through the use of replica exchange simulations, characterise (at least the initial stages) of conformational change at oil-water interfaces.

**FIG. 1.**
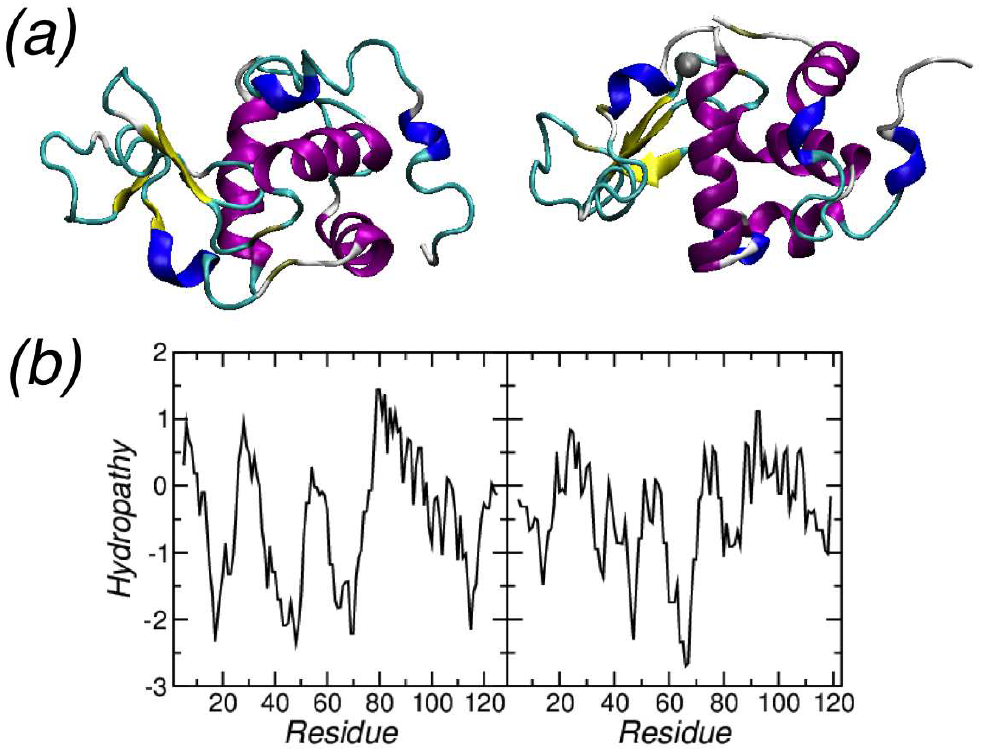
(a) Structures of LSZ (left) and *α*La (right). (b) Kyte-Doolite hydropathy plots for LSZ (left) and *α*La (right). Generated using the ExPASy webserver^37^ (http://web.expasy.org/cgi-bin/protscale/protscale.pl)

## II. SIMULATION DETAILS

### A. Simulated systems

The simulated systems consisted of a single protein, either lysozyme or *α*-lactalbumin, and a preequilibrated water-octane interface. Initial protein structures were taken from the protein database, 1E8L for LSZ^38^ and 1HFZ for *α*-lactalbumin^39^. For *α*-lactalbumin the experimental structure contained a V90M mutation; this was reversed in VMD^40^ using the psfgen plugin. Charges on polarizable residues were set appropriately for pH=7. Chloride or sodium counter ions were added as needed to neutralise the system. The proteins were placed in the water slab, approximately 40 Å from the interface. For each protein three starting orientations were used (shown in Figure 2). In the first the protein principal axes were orientated along the *x*, *y*, and *z* axes, with the long axis along the *x*-axis and the short along the *z*-axis. In the second the protein long axis was still orientated along the *x*-axis but the protein short axis was orientated along the *y*-axis. In the final orientation the protein long axis was orientated along the *z*-axis and the short along the *y*-axis. Simulations from each of the different starting orientations will denoted as Rotx, Roty, and Rotz in the rest of this paper.

**FIG. 2.**
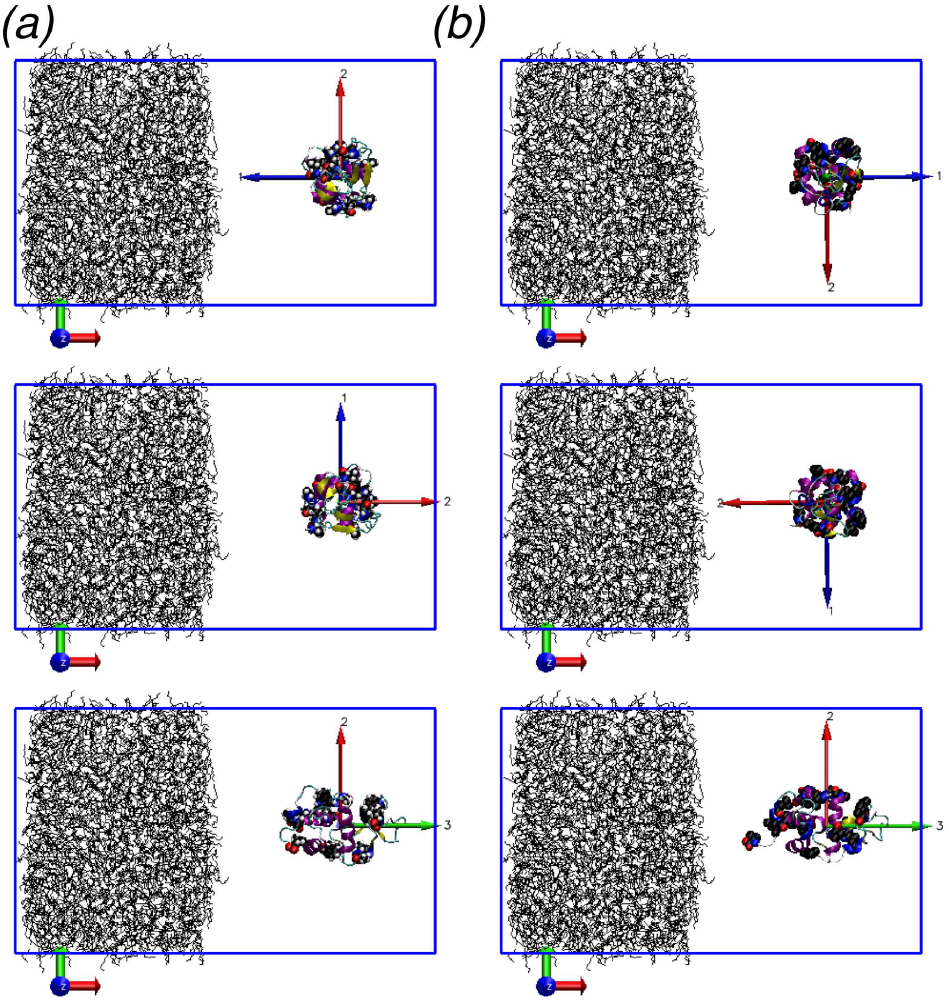
Initial configurations for (a) LSZ and (b) *α*-lactalbumin simulations. Long axis orientated along *x*, *y*, and *z* directions (top to bottom). Exposed hydrophobic residues (with solvent accessible surface areas greater than 30 Å^2^ in crystal structure) highlighted by van der Waals spheres. Water omitted for clarity.

### B. Methodology

The simulations were performed using the Gromacs molecular dynamics package^41,42^. For the REST simulations version 4.6.3 using a modified version of PLUMED^43,44^ was employed, with Gromacs 5.0 used for the other simulations. Standard gromacs tools were used to create the simulation input files. The Gromos54a7 force field^45^ was used for modelling the proteins and along with SPC water^46^. The Gromos family of force fields have been parameterised against solvation free energies for amino acid side chains^45,47^ so may be expected to give a good description of the partitioning of these between water and hydrophobic media and have been used in a number of previous studies of proteins at liquid interfaces^20,21,33,48,49^.

Following previous work^20,21^ van der Waals and real-space electrostatic interactions were truncated at 10 Å. Long-range electrostatic interactions were accounted for using PME^50^ with a reciprocal space grid of 56 × 56 × 96

The simulations were performed in the *NpT*-ensemble with the temperature controlled using a velocity rescaling algorithm^51^ (relaxation time 0.1 ps) and pressure was controlled using the Parrinello-Rahman barostat, with relaxation time 2 ps. Bond lengths were constrained using the LINCS algorithm^52^. All systems were energy minimized using the steepest descent algorithm, followed by a short *NVT* simulation (20 ps) with the heavy atoms in the proteins restrained to their initial positions using harmonic potentials with a force constant of 1000 kJ mol^−1^ nm^−2^. Following this short *NVT* and *NpT* simulations (both 20 ps), without restraining the heavy atoms, were performed.

Two sets of simulations were performed for each protein. For the three different starting conformations a standard *NpT* MD simulation, with the protein starting in the water region, were performed. These simulations were used to investigate the initial interfacial adsorption and its dependence on the protein orientation. Following this replica exchange with solute tempering (REST) simulations^44,53^ were performed to improve sampling over different protein conformations. This is a variation on replica exchange molecular dynamics^54^ where the temperature of only a subset of the system, in this case the protein, varies between replicas. This allows for the use of a smaller number of replicas compared to standard REMD. The temperature scaling is effected by scaling the protein-protein and protein-system interactions; the potential energy is given by

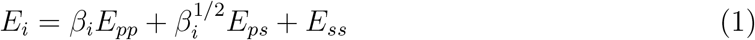

where *E*_*pp*_, *E*_*ps*_, and *E*_*ss*_ are the protein-protein, protein-solvent (both water and octane), and solvent-solvent interaction and the scaling factor *β*_*i*_ = *T*_0_*/T*_*i*_. In these simulations 8 replicas were used with temperatures (scaling factors) of 298 K (1.0), 307.71 K (0.968), 317.74 K (0.938), 328.09 K (0.908), 338.79 K (0.880), 349.83 K (0.852), 361.23 K (0.825), and 373 K (0.80). *NpT* simulations performed at the highest temperature showed that this was sufficient for both proteins to unfold. For both LSZ and *α*La two REST simulations, starting from the Rotx and Rotz *NpT* simulations, were performed. From the *NpT* simulations these had the most and fewest residues in contact with the interface. All discussion of results of the REST simulations (Sections IV-VI) refer to the *β*_0_ = 1 (298 K) replica as this is the only physically relevant case.

The standard *NpT* and REST simulations were run for 500 ns and 400 ns respectively, with coordinates were saved every 10 ps for later analysis. Prior to the REST simulations *NpT* simulations of up to 500 ns were performed at each of these temperatures, starting from the end of the 298 K *NpT* simulation in order to equilibrate the system at these higher temperatures. These higher temperature simulations were ran until the protein structure, measured through the radius of gyration, had reached a constant value.

### C. Analysis

The simulations were analysed using a combination of custom written python scripts (using the MDAnalysis library^55^), VMD^40^, and standard Gromacs utilities. Simulation snapshots were generated using VMD, with the protein secondary structure determined using the STRIDE algorithm^56^. Protein conformations were clustered using the Gromacs *g_cluster* utility according to the Gromos criteria^57^, with *R*_*cut*_ = 3 Å. The potential energy contributions were calculated using the rerun option of *mdrun*.

The position of the oil-water interface was determined from the Gibbs dividing surfaces for the water 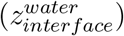 and octane 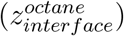 as 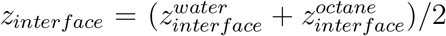. The location of the dividing surfaces for the water and octane were found following Vink *et al*^58^. To estimate the surface area, a grid of points is placed on the water-octane interface, defined by the Gibbs surface with the area being estimated from the number of grid points that were within a cutoff distance from a protein atom, given by its van der Waals radius+1.4 Å (van der Waals radius of water oxygen). The partition free energy was estimated by summing the water-oil transfer free energies for each residue whose side-chain centre of mass is in the octane (*z < z*_*interface*_), using the water-cyclohexane transfer free energies of the amino acid side chains^59^. In the calculation of the partition free energy only residues that were solvent accessible in solution (taken to be those with a solvent accessible area above 30 Å^2^ in the experimental structure) were considered as other residues would not be expected to undergo a significant change in energy on going from the hydrophobic core to the apolar oil phase. While experimental values of the water-octane transfer free energies for amino acid side chains are not known the chemical similarity between cyclohexane and octane suggests that this is reasonable approximation.

Following previous work the number of hydrophobic contacts and the number of protein-octane contacts were determined using^22^

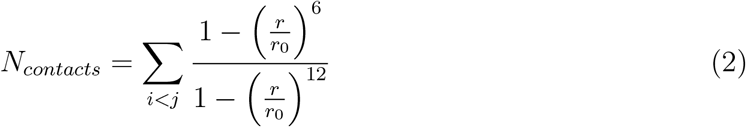

where *r*_0_ = 5 Å. For hydrophobic contacts within the protein the sum runs over the C_*γ*_ atoms of the hydrophobic residues (for alanine the *C*_*β*_ atom is used). For protein-octane contacts the sum runs over C_*γ*_ atoms of the hydrophobic residues and all the united carbon atoms in the octane molecules, with the results for each octane molecule divided by eight. The switching function in Eqn 2 is defined so that it goes to 1 at small *r* and 0 at large *r*.

## III. LYSOZYME AND *α*-LACTALBUMIN ADSORB TO WATER-OCTANE INTERFACE IN DIFFERENT MANNERS

Shown in Figure 3(a) is the protein centre-of-mass-interface separation for LSZ. Independent of starting orientation the protein adsorbs onto the interface within ~ 20 ns. Examination of the separations of individual residues and the interface shows that these differ markedly for the three simulation runs (Figure 3(b), suggesting the adsorption is essentially non-specific. These typically include clusters of residues around surface exposed hydrophobic residues. This leads to a number of hydrophilic amino acids being drawn into the oil phase. Simulations started from the Rotx and Roty orientations show similar numbers of residues in contact with the interface; these residues include both the N and C-termini of the protein. Between these the residues in contact with the interface differ between these two simulations (Figure 3(c)). For the Rotx simulation a cluster of residues around the hydrophobic F34 and F38 residues move into the oil. This region is more distant from the interface in the Roty simulation. Instead the section G102-N106 enters the octane. The Rotz simulation has fewer residues in contact with the interface. These are also quite distinct from those for Rotx and Roty with contact made with the interface between residues N77 and S85. In all cases the residues in contact with the interface correspond to hydrophobic regions of the protein, shown by peaks in the hydropathy plot for LSZ (Figure 1(b)).

**FIG. 3.**
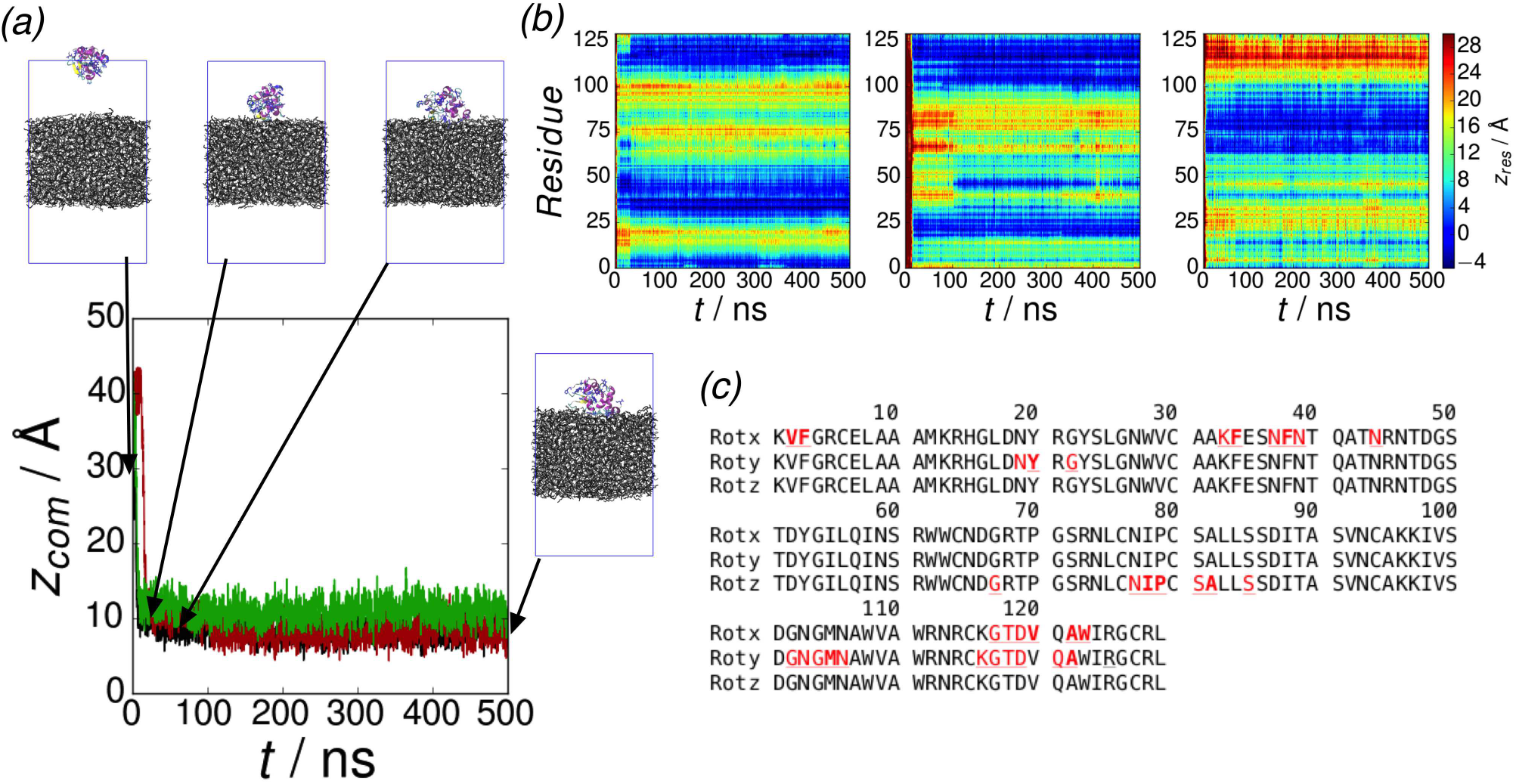
Approach of LSZ to water-octane interface. (a) Protein centre-of-mass-interface separation. Rotx, Roty, and Rotz starting configurations denoted by black, red, and green lines respectively. Insets show simulation snapshots for Rotx at *t* =0, *t* = 10 ns, *t* = 50 ns, and *t* = 500 ns (b) Residue centre-of-mass-interface separations for (left to right) Rotx, Roty, and Rotz starting configurations. (c) Annotated sequence with residues in the octane slab (*z*_*res*_ < 0 for the final 100 ns) underlined. Hydrophobic residues in octane slab shown in bold.

Unlike LSZ *α*La can show a significant lag time for interfacial adsorption (Figure 4(a)). Depending on the starting orientation attachment to the interface occurs after ~ 40 − 80 ns. Examination of the individual residue separations (Figure 4(b)) suggests that this is associated with reorientation of the protein from its initial orientation to one which favours interfacial adsorption. For all the starting orientations similar sets of residues are in contact with the interface (Figure 4(c)). This similarity between the residues involved in interfacial attachment suggests that this is more strongly dependent on the protein than the nonspecific adsorption seen for lysozyme. *α*La contains two amphiphilic helices, helix A (E1-L15) and helix C (T86-V99) that mediate its binding onto membranes^60^; from analysis of the simulations it can be seen that helix C is also involved in the adsorption of *α*La onto the water-octane interface, with a region between helices A and B (L23-S34)^39^ also involved. The initial reorientation of the protein can be seen in the simulation snapshots (Figure 4(a)). At short times the protein reorients near the centre of the water slab until a possible binding orientation is found. Once it is oriented correctly attachment to the interface occurs over similar timescales as for LSZ. The final adsorbed orientation and structure is similar for all the starting orientations. Neither LSZ or *α*La exhibit transient interfacial attachment observed in molecular dynamics simulations of other proteins^20,22^.

**FIG. 4.**
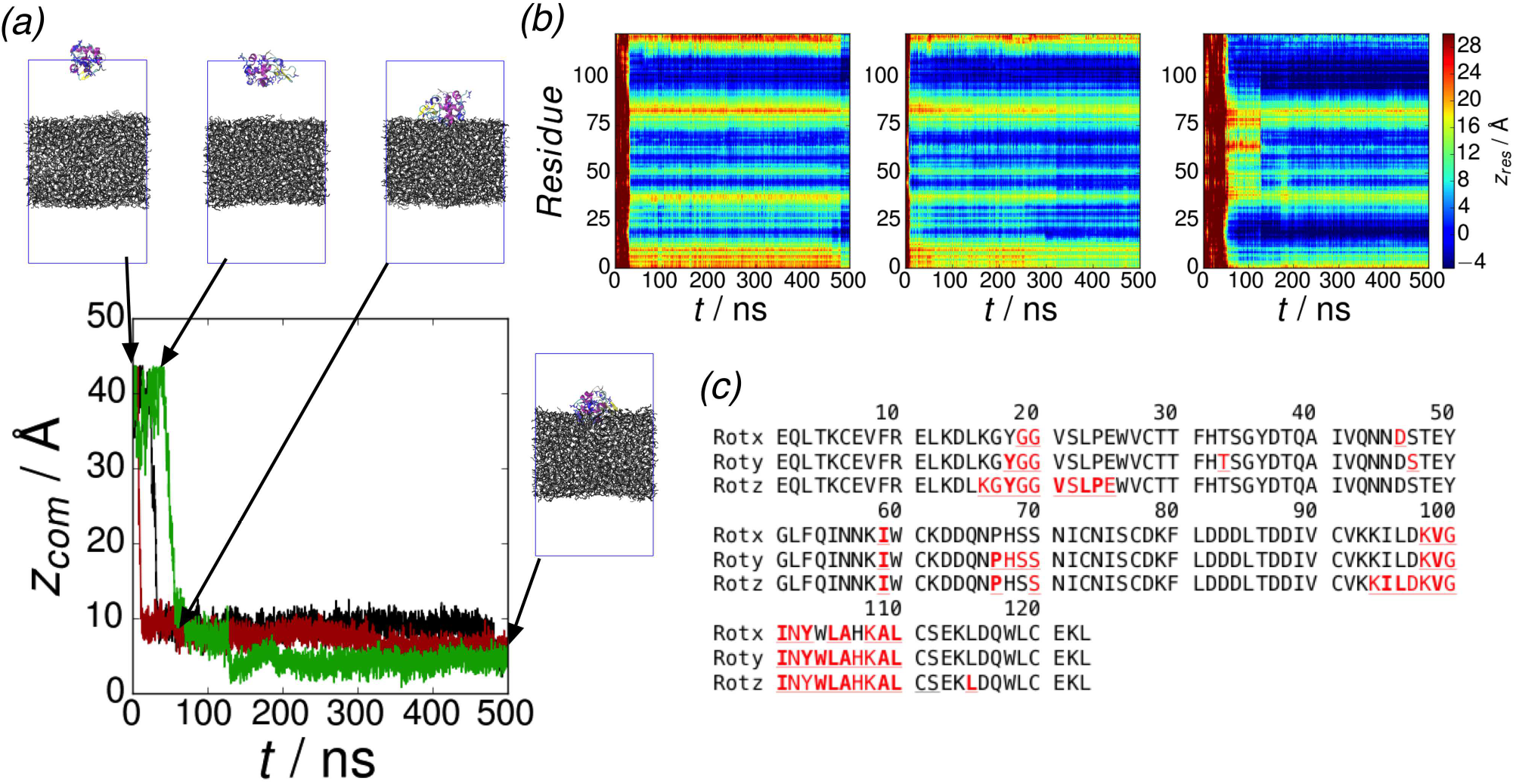
Approach of *α*La to water-octane interface. (a) Protein centre-of-mass-interface separation. Rotx, Roty, and Rotz starting configurations denoted by black, red, and green lines respectively. Insets show simulation snapshots for Rotx at *t* =0, *t* = 25 ns, *t* = 50 ns, and *t* = 500 ns (b) Residue centre-of-mass-interface separations for (left to right) Rotx, Roty, and Rotz starting configurations. (c) Annotated sequence with residues in the octane slab (*z*_*res*_ < 0 for the final 100 ns) underlined. Hydrophobic residues in octane slab shown in bold.

During the adsorption only slight changes in the secondary structure are seen for LSZ (Figure 5(a)). In some of the simulation runs (Rotx and Rotz) there is some loss of *α*-helix content towards the C-terminal end of the protein (residues 120-125). From the residue centre-of-mass positions (Figure 3(b)) there is no relationship between this and the location of these regions relative to the interface.

**FIG. 5.**
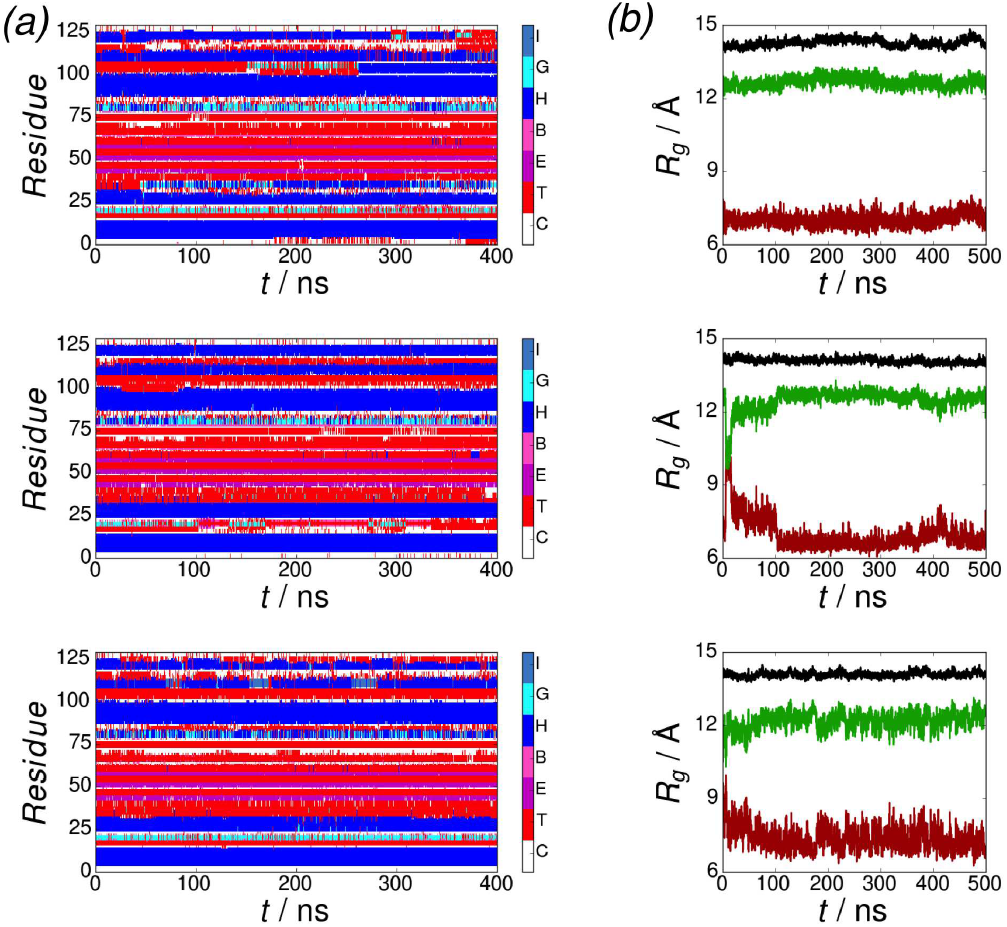
Protein structure during interfacial adsorption for LSZ at 298 K. (a) Secondary structure for (top to bottom) Rotx, Roty, and Rotz simulations. Blue denotes *α*-helix, cyan 3_10_-helix, red turn, pink bridge, magenta extended, and white random coil. (b) Radius of gyration for (top to bottom) Rotx, Roty, and Rotz simulations. *R*_*g*_, 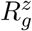, and 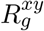 denoted by black, red, and green lines respectively.

Similarly little change is seen in the radius of gyration (Figure 5(b)). For all simulations the overall radius of gyration remains essentially unchanged suggesting that protein shape doesn’t change during adsorption. The perpendicular and in plane components of the radius of gyration show some change, consistent with protein reorientation during adsorption.

For *α*La more variation is seen in the secondary structure (Figure 6(a)). While the membrane binding helices A and C are largely unchanged during this initial adsorption the other helices change size during this. Helix D (A106-C111) shows the largest change, with this shrinking or disappearing altogether. The unstable nature of helix D may be expected, as in bulk solution this helix is found to be pH-dependent. This helix lies in a region that is closer to the interface, with a substantial portion of this penetrating into the oil-phase. Compared to helices A and C it is less amphiphililic so this penetration into the oil phase distorts this helix. Using a helical wheel analysis^61^ the hydrophobic dipoles for helices A, C, and D are 4.44, 1.51, and 1.09 (calculated using the Wimley-White interface hydrophobicity scale^62^) respectively.

**FIG. 6.**
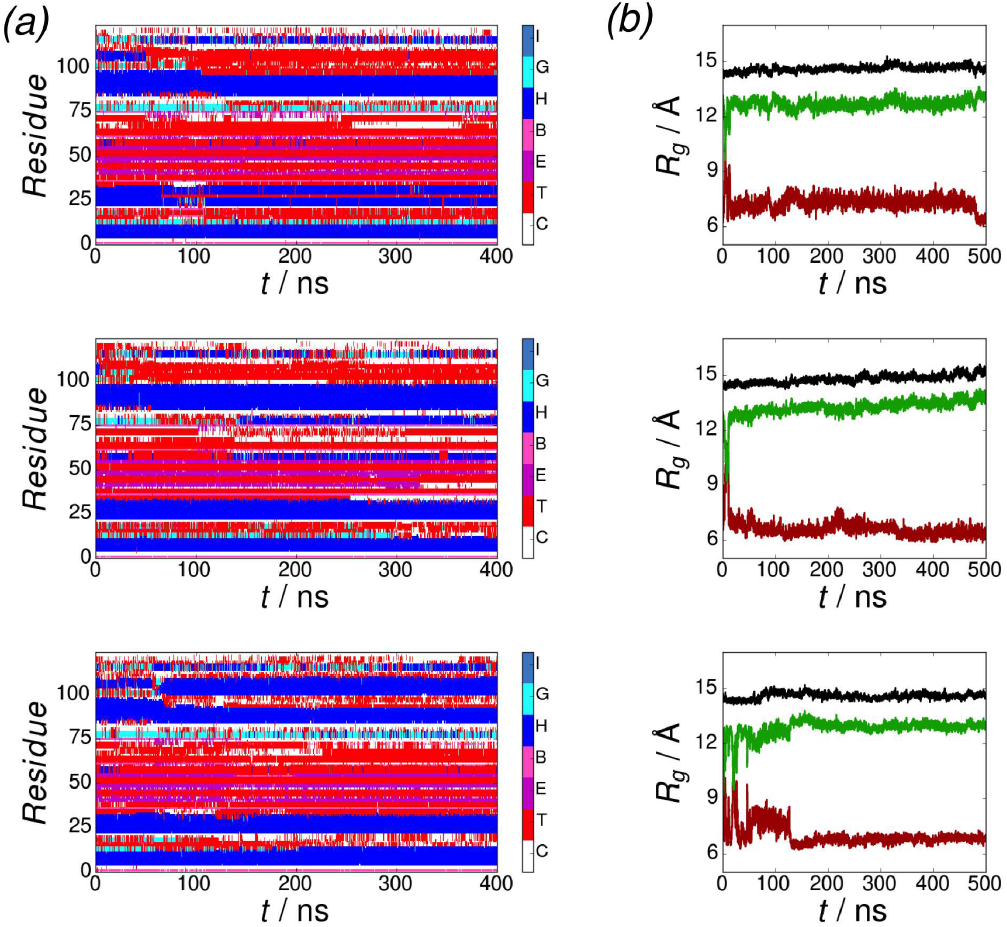
Protein structure during interfacial adsorption for *α*La at 298 K. (a) Secondary structure for (top to bottom) Rotx, Roty, and Rotz simulations. Blue denotes *α*-helix (H), cyan 3_10_-helix (G), red turn (T), pink bridge (B), magenta extended (E), and white random coil (C). (b) Radius of gyration for (top to bottom) Rotx, Roty, and Rotz simulations. *R*_*g*_, 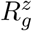, and 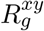 denoted by black, red, and green lines respectively.

Similar to LSZ the tertiary structure shows little change during interfacial adsorption (Figure 6(b)). The overall *R*_*g*_ is largely constant during adsorption, while 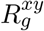 and 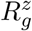 change consistent with the protein reorienting during adsorption.

## IV. PROTEIN CONFORMATIONAL CHANGE AT WATER-OCTANE INTERFACE

The REST simulations reveal that both LSZ and *α*La exhibit changes in secondary and tertiary structure at the octane-water interface. For LSZ the radius of gyration, both overall and the components resolved perpendicular and in the plane of the interface, shows rapid jumps between two sets of values (Figure 7(a)). This suggests that LSZ can adopt (at least) two different structures at the interface. For one of these the radius of gyration is similar to the value for bulk solution suggesting that this corresponds to a compact, globular structure similar to the solution state. The second has significantly larger 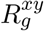 and smaller 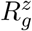 showing that the protein has expanded in the plane of the interface. For both simulations the compact state is significantly more common. This is consistent with simulations of LSZ at the water-DCE interface^32^, which also found that LSZ preferentially adopts a compact state but more expanded conformations were observed during the simulations. It is also similar to recent simulations of myoglobin-derived peptides, where both compact and globular states were found^33^. The lack of significant tertiary structure change is also seen in the locations of the residues (Figure 7(b)) which are essentially the same as for the *NpT* simulations.

**FIG. 7.**
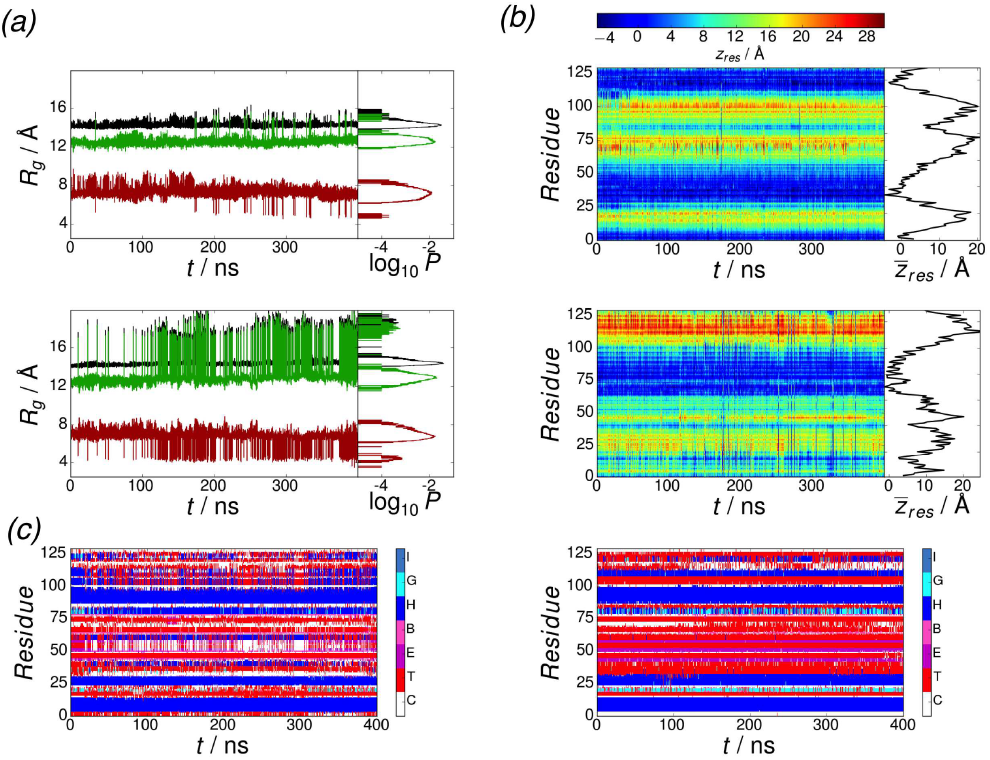
Structure of LSZ at 298 K from REST simulations. (a) Radius of gyration for Rotx (top) and Rotz (bottom) simulations. Black, red, and green lines denote *R*_*g*_, 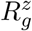, and 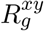 respectively. Right hand panels show probability histograms for *R*_*g*_, 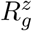, and 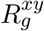 (symbols as in main panel) calculated over final 100 ns of each simulation. (b) Residue-interface separations for Rotx (top) and Rotz (bottom) simulations. Right hand panels show average residue-interface separations calculated over final 100 ns of simulations. (c) Secondary structure distributions for Rotx (left) and Rotz (right) simulations. Blue denotes *α*-helix (H), cyan 3_10_-helix (G), red turn (T), pink bridge (B), magenta extended (E), and white random coil (C).

These different conformations also exhibit slightly different secondary structure compositions. While the compact state has similar secondary structure composition to the solution state, the *α*-helix content of the expanded state is slightly lower (Table I). This is particularly noticeable for the C-terminal end of the protein, where these helical regions typically become turns. Despite the similarity between the secondary and tertiary structures between the two REST simulations different sections of the protein are in contact with the interface. These regions are the same as in the *NpT* simulations, indicating that we do not see reorientation of the protein. Again this suggests that for LSZ adsorption to the water-octane interface is non-specific.

**TABLE I.**
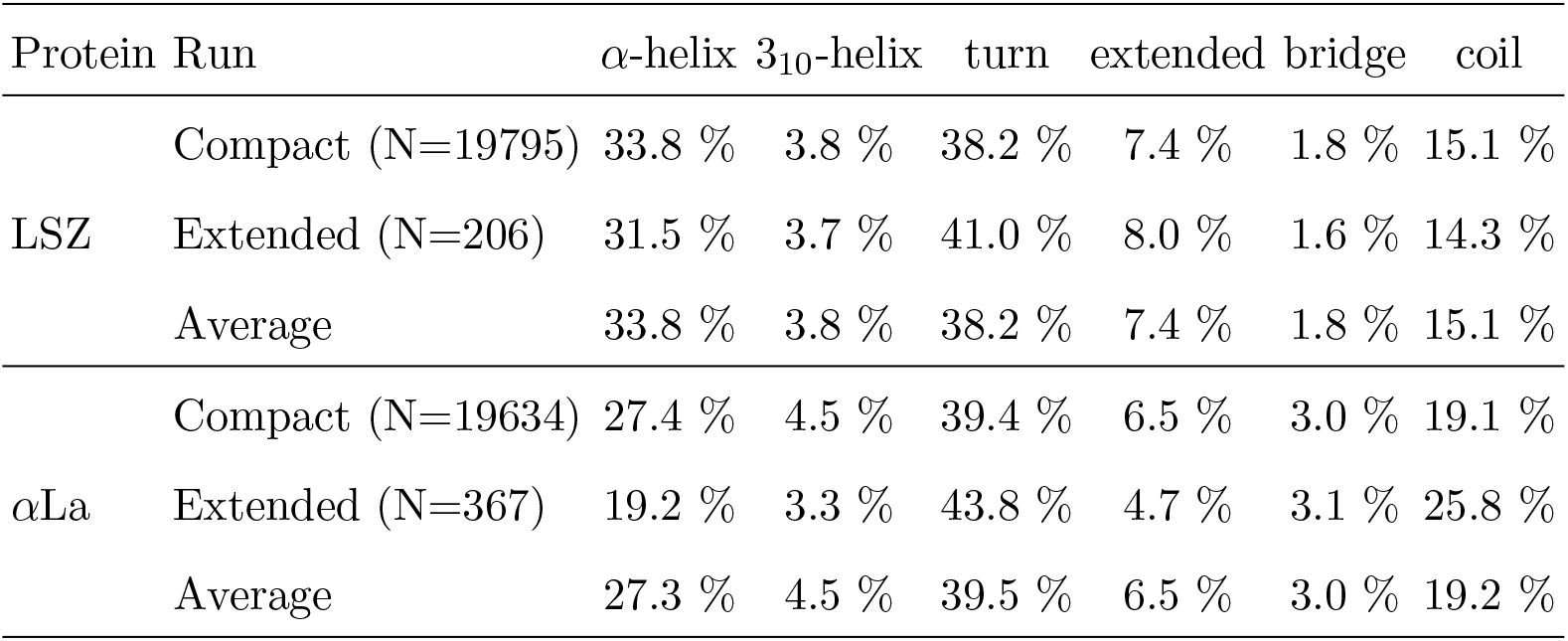
Secondary structure composition from REST simulations decomposed into compact and extended states.

| Protein | Run | α-helix | $3_{10}$-helix | turn | extended | bridge | coil |
| --- | --- | --- | --- | --- | --- | --- | --- |
| LSZ | Compact (N=19795) | 33.8 % | 3.8 % | 38.2 % | 7.4 % | 1.8 % | 15.1 % |
|  | Extended (N=206) | 31.5 % | 3.7 % | 41.0 % | 8.0 % | 1.6 % | 14.3 % |
|  | Average | 33.8 % | 3.8 % | 38.2 % | 7.4 % | 1.8 % | 15.1 % |
| αLa | Compact (N=19634) | 27.4 % | 4.5 % | 39.4 % | 6.5 % | 3.0 % | 19.1 % |
|  | Extended (N=367) | 19.2 % | 3.3 % | 43.8 % | 4.7 % | 3.1 % | 25.8 % |
|  | Average | 27.3 % | 4.5 % | 39.5 % | 6.5 % | 3.0 % | 19.2 % |

While both simulation runs for *α*La started from similar conformations, the protein structures found in these simulations are noticeably different (Figure 8). Starting from the Rotx conformation only a single compact state is found, while both compact and extended states are found from the Rotz conformation (Figure 8(a)). Compared to LSZ the extended state for *α*La is smaller (*R*_*G*_ ~ 17 Å compared to ~ 20 Å for LSZ). For the Rotz simulation these changes in structure can be seen in the residue centre-of-mass positions (Figure 8(b)). In the Rotz simulation helix-B becomes destabilised and some changes to the *α*-helix content towards the C-terminus are seen in both simulations. The residue-interface separations for these regions differ between the two simulation runs with loss or destabilisation of *α*-helical regions being more common when these are closer to the interface. Helix D is less amphiphilic than A and C that are largely unchanged at the interface, so when this is becomes close to the interface the helices will become distorted to allow hydrophobic side chains to partition into the oil. Considering the compact and extended conformations for *α*La the secondary structures of these show a larger difference than LSZ. In particular the extended states show a much smaller proportion of *α*-helix; compact states have ≈ 27 % while extended states this drops to ≈ 19 %, with the turn and random coil proportion increasing.

**FIG. 8.**
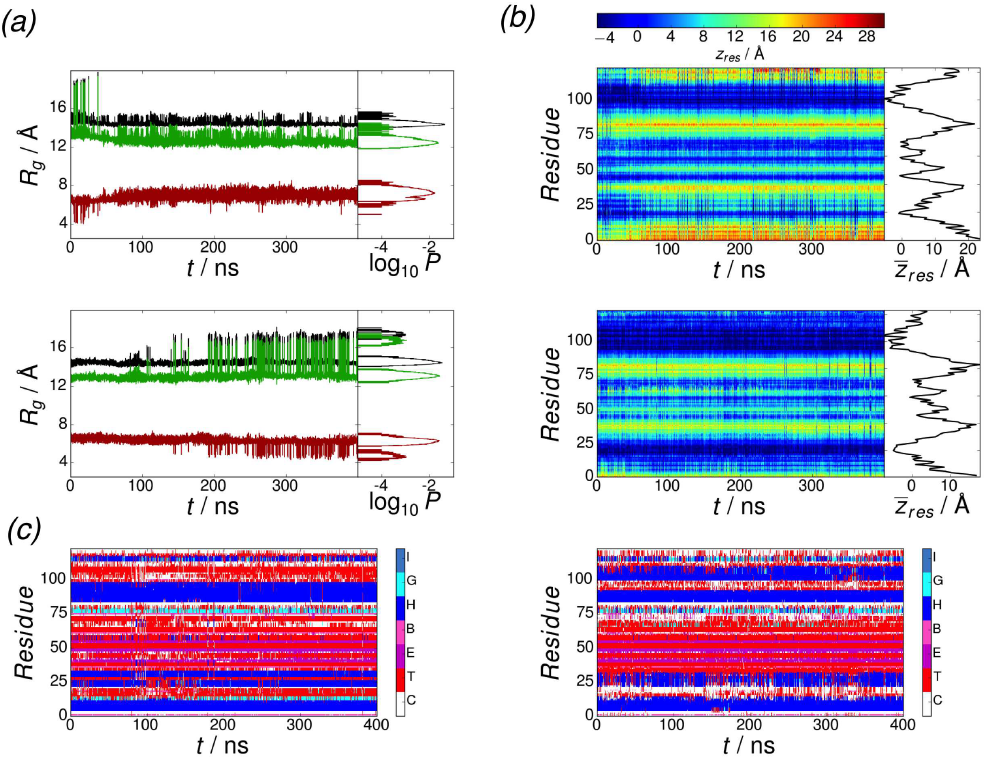
Structure of *α*Laat 298 K from REST simulations. (a) Radius of gyration for Rotx (top) and Rotz (bottom) simulations. Black, red, and green lines denote *R*_*g*_, 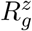, and 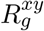 respectively. Right hand panels show probability histograms for *R*_*g*_, 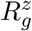, and 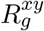 (symbols as in main panel) calculated over final 100 ns of each simulation. (b) Residue-interface separations for Rotx (top) and Rotz (bottom) simulations. Right hand panels show average residue-interface separations calculated over final 100 ns of simulations. (c) Secondary structure distributions for Rotx (left) and Rotz (right) simulations. Blue denotes *α*-helix (H), cyan 3_10_-helix (G), red turn (T), pink bridge (B), magenta extended (E), and white random coil (C).

## V. LYSOZYME AND *α*-LACTALBUMIN EXHIBIT MULTIPLE INTERFACIAL CONFORMATIONS

Analysis of the radius of gyration suggests that both LSZ and *α*La exhibit multiple conformations at the oil-water interface. For both proteins compact, globular states are more common, with more extended states being rarer. Both LSZ and *α*La possess a network of disulfide bonds helping them retain their structure. *α*La is additionally stabilised by its Ca^2+^ ion. These strong interactions make them resistant to changes in tertiary structure at interfaces, unlike the recently studied case of myoglobin-derived peptides that lack these.

To get more insight into the different protein conformations we performed a cluster analysis on the last 100 ns of each of the REST simulations for both proteins (20000 protein conformations in total). For LSZ 17 distinct clusters were found but only one of these is significantly occupied (Figure 9(a)). This corresponds to the centre of the cluster of compact states (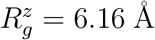 and 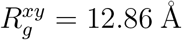) in the probability map (Figure). From the radius of gyration of each cluster the first six clusters are all relatively compact. A number of distinct extended states were found, however, the probability of these are significantly lower than the compact states. We can estimate the free energy of each cluster from ∆*F*_*i*_ = −*k*_B_*T*ln *P_i_/P*_1_ where *P*_*i*_ is the probability of the protein being in the *i*th cluster. There is a large jump in free energy difference the first and second clusters, illustrating the dominance of the first cluster. The extended states have ∆*F*_*i*_ ≳ 3 kcal mol^−1^.

**FIG. 9.**
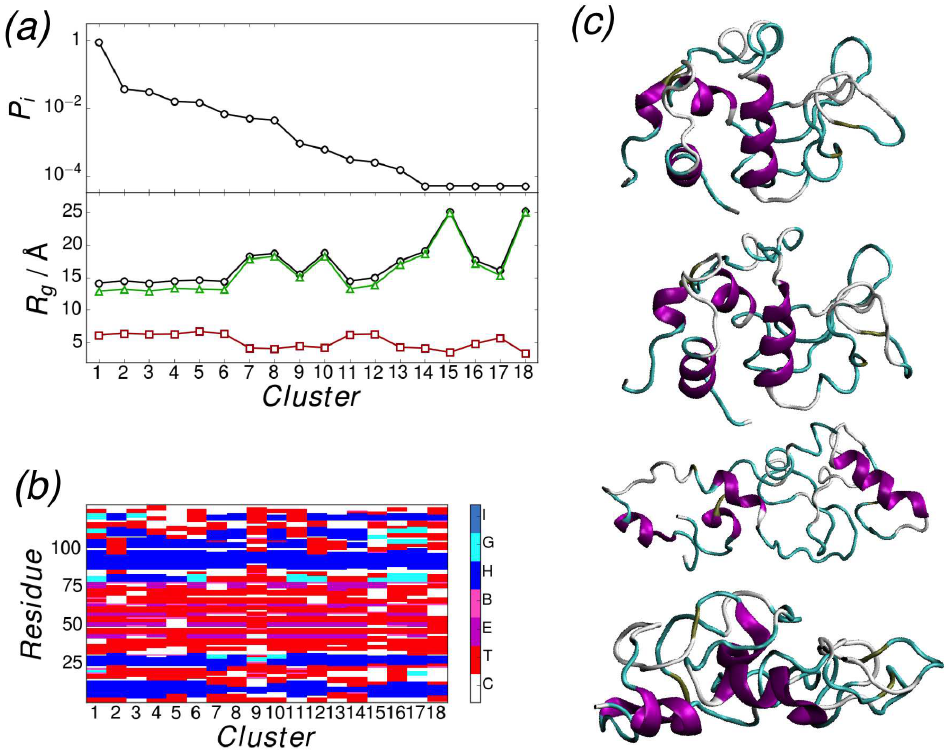
(a) Cluster probability (top) and radius of gyration (bottom) for each of the cluster average conformations for LSZ. In bottom panel *R*_*g*_, 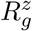, and 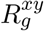 denoted by black, red, and green lines respectively. (b) Secondary structure for the cluster average structures. (c) Structures of first, second, seventh, and ninth (top to bottom) cluster average structures.

Despite the differences in tertiary structure the secondary structure is largely the same for the different conformations (Figure 9(b)). With the exception of one cluster, most variation is seen towards the C-terminus, from residue G102 onwards. Most commonly there is a short *α*-helical region (G102-A107) followed by regions of turn and 3_10_-helix, as in the first cluster. In the conformations this helix is absent and other *α*-helix regions are found (e.g. A110-C115 and V120-I124 in the second cluster). Interestingly this holds for most of the extended conformations, suggesting that for the most part these consist of different arrangements of the helical regions. The only cluster that deviates significantly from this is the 9th which has only two *α*-helical regions (E7-A11 and V92-V99). This corresponds to the small cluster of states between the compact and extended states (Figure 15). Snapshots of the cluster average structures (Figure 9(c)) show that the most common compact conformations (for example the 1st and 2nd conformations) are similar to each other, whereas more differences are seen for the more extended states. The 7th cluster, which is the first extended conformation, has a significantly smaller amount of *α*-helix and is highly extended in the plane of the interface. For the 9th cluster, which has the lowest *α*-helix content, the protein adopts a compact, though elongated structure that is quite different to the other clusters.

In the cluster analysis for *α*La only eight distinct conformations were found (Figure 10(a)). The first two are both compact structures and are approximately equally occupied, consistent with the probability map. There is a smaller free energy difference between compact and extended states for *α*La (∆*F*_*i*_ ~ 1.9 kcal mol^−1^).

**FIG. 10.**
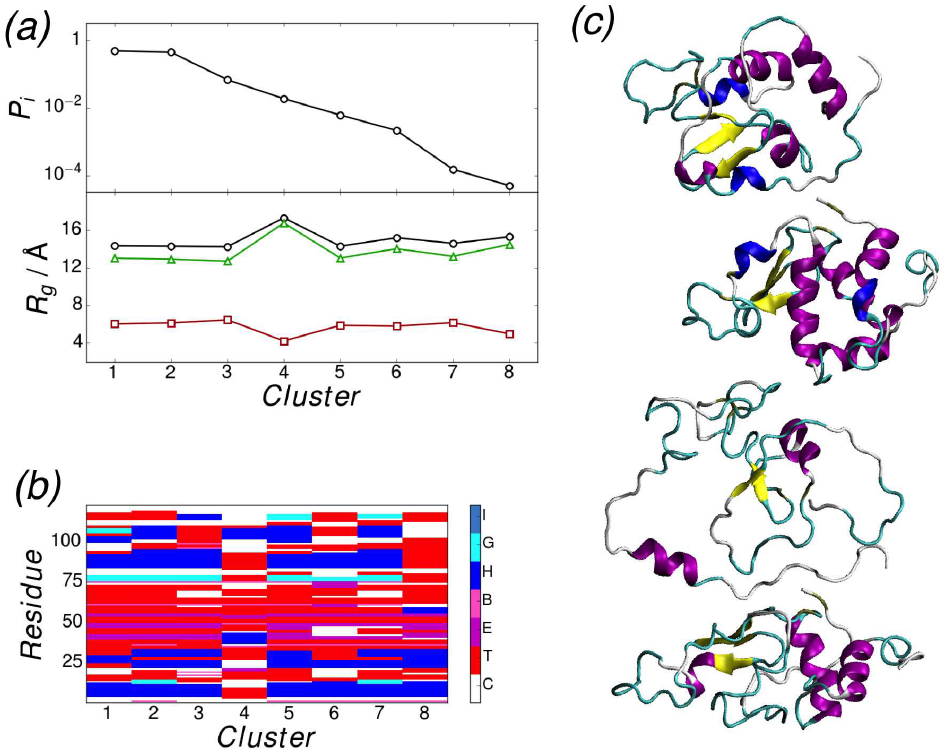
(a) Cluster probability (top) and radius of gyration (bottom) for each of the cluster average conformations for *α*La. In bottom panel *R*_*g*_, 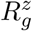, and 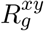 denoted by black, red, and green lines respectively. (b) Secondary structure for the cluster average structures. (c) Structures of first, second and fourth (top to bottom) cluster average structures.

Compared to LSZ more variation in seen the secondary structures for the different clusters (Figure 10(b)). As is seen in the time series of the secondary structure the membrane-binding helices (A and C) are present in almost all the clusters. The other helices are more variable, with one or other of these missing in some of the clusters. Only in the extended conformations (clusters 4 and 8) are major differences in these areas seen. For cluster 4 neither of the membrane-binding helices are present. Instead helix-D is largely present (Y103-A109) and an additional short *α*-helix (D38-N44) segment is formed.

Snapshots of these different clusters show clearly how the compact and extended structures differ from each other (Figure 10(c)). While the compact structures are quite similar to each other, the first extended structure is significantly different. It is highly extended in the plane of the interface extending with both *α*-helices lying flat against the water-octane interface. The second, less common, extended structure is more closers to the compact conformations.

## VI. WHAT FACTORS CONTROL CONFORMATIONAL CHANGE?

Changes in protein conformation at water-oil interfaces are driven by a number of effects, including interfacial free energies, partitioning of hydrophobic residues, and differing protein-solvent interactions between the two different solvent components. To gain insight into these different effects we can estimate the differences between these for the extended and compact states, which reveals some differences between the interfacial behaviour of LSZ and *α*La. The free energy difference between different interfacial conformations can be estimated from

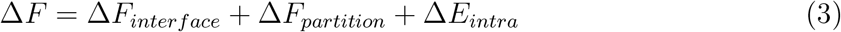

where the terms on the right hand side are the changes in interfacial free energy, partition free energy, and protein intramolecular energy respectively. The interfacial free energy is given by −*γA*_*o*_, where *γ* is the water-octane interfacial tension and *A*_*o*_ is the area occupied by protein (details given in methodology section). *γ* was calculated from molecular dynamics simulations of a water-octane interface, in the absence of a protein, using

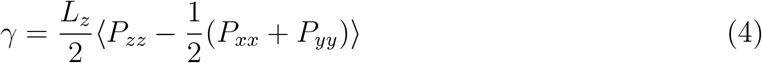

which gave a value of 51.8 mN m^−1^, in good agreement with the experimental value^63^ (*γ* = 51.2 mN m^−1^). Estimates for these different contributions are presented in Table II, along with the protein-octane (*E*_*po*_) and protein-water interaction (*E*_*pw*_) energies.

**TABLE II.** Contributions to energies of compact and extended states for LSZ and *α*La. For both proteins conformations with *R*_*g*_ ≤ 16 Å were considered compact and *R*_*g*_ > 16 Å were considered extended. Averages calculated over final 100 ns of each REST simulation. *A*_*o*_ and *γ* are the occupied surface area and the water-octane interfacial tension .Areas in Å^2^, energies in kcal mol^−1^. Uncertainties estimated as standard errors.

| Protein | Conformation | $N$ | $A_o$ | $-\gamma A_o$ | $\Delta F_{partition}$ | $E_{intra}$ | $E_{po}$ | $E_{pw}$ |
| --- | --- | --- | --- | --- | --- | --- | --- | --- |
| LSZ | Compact | 1981 | 269±2 | -20.1±0.15 | 25.8±0.6 | -3218±1.5 | -114.7±0.8 | -1806±2 |
| | Expanded | 21 | 740±12 | -54.8±0.9 | 30±4 | -3100±20 | -120±15 | -1830±16 |
| αLa | Compact | 1952 | 366±2 | -27.3±0.15 | 11.4±0.3 | -3737±3 | -166.3±0.9 | -2107±3 |
| | Expanded | 50 | 676±8 | -50.4±0.6 | 16±2.5 | -3590±12 | -290±1.6 | -2060±20 |

For both proteins the interfacial free energy favours extended states at the interface, as may be expected. The extended states occupy a larger area at the interface, giving rise to a lower interfacial free energy compared to the compact state. In the extended states the hydrophobic residues are able to partition more easily into the oil phase decreasing the partition free energy. The partition free energy undergoes a slight increase for both LSZ and *α*La, due to the unfavourable partitioning of polar sidechains into the octane. The change in *F*_*interface*_ on going from compact to extended states is larger for LSZ than *α*La, while the change in *F*_*partition*_ are approximately the same for both proteins.

While the interfacial free energy favours extended states for both proteins the intramolecular energy prefers the compact states. The contributions from the protein-solvent interactions behave differently for the two proteins, most significantly the protein-octane interaction. For *α*La this is significantly lower for the extended state compared to the compact state, whereas for LSZ this is essentially unchanged. This suggests that the conformational change for *α*La is driven by favourable interactions between the protein and oil molecules. The protein-octane interaction in the compact state for *α*La is more favourable than LSZ, consistent with the lower partition free energy. The protein-water interaction only changes slightly for both proteins; for LSZ this decreases slightly going from compact to extended states, while it increases for *α*La.

To understand the relationship between the intramolecular and protein-octane interaction energies and the protein conformation we can examine the probability distributions *P*(*R*_*g*_, *E*_*intra*_) and *P*(*R*_*g*_, *E*_*po*_) (Figure 11). For both LSZ and *α*La *P*(*R*_*g*_, *E*_*intra*_) is similar for the compact states, with a narrow distribution of *R*_*g*_ values and the *E*_*intra*_ values spanning a range of a few 100 kcal mol^−1^. For LSZ these take on a wider range of *R*_*g*_ values compared to *α*La; in both cases they span a similar range of energies as in the compact states but are skewed towards higher energies, reflecting the weaker intramolecular interactions for the extended states.

**FIG. 11.**
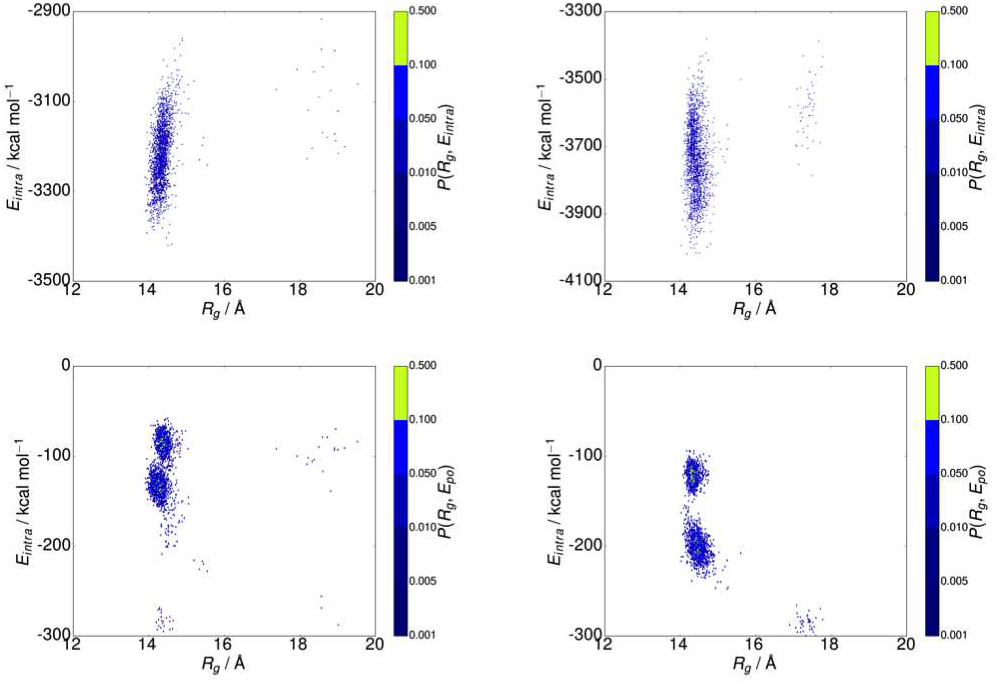
Probability maps for radius of gyration and protein intramolecular interaction (top) and protein-octane interaction (bottom) for LSZ (left) and *α*La (right)

More variation is seen in *P*(*R*_*g*_, *E*_*po*_). For *α*La almost all states are concentrated into three distinct groups, two at lower *R*_*g*_, corresponding to compact states, and a third extended group. The most extended group has a lower *E*_*po*_ than the compact ones showing, as would be intuitively expected, that the extended state allows for more favourable interactions between protein and oil. LSZ shows more groups, consistent with the higher number of conformations found in the cluster analysis. It possesses a number of distinct compact conformations with different protein-octane interaction energy. The extended states typically do not have a lower *E*_*po*_ than the compact states. This shows that, unlike *α*La, conformational change of LSZ at the water-octane interface is not, at least in its initial stages, driven by favourable interactions between the protein and oil.

When the proteins go from compact to extended states this changes the number of contacts between hydrophobic residues and introduces contacts between octane molecules and hydrophobic residues that normally form the compact protein core (Figure 12). Both of these show bimodal behaviour with frequent transitions between values appropriate for compact and extended states. The number of contacts between hydrophobic residues in the compact states is similar for both LSZ and *α*La, while this is lower for LSZ in the extended states. *α*La typically has more contacts between hydrophobic residues and octane, consistent with the more favourable *E*_*po*_ compared to LSZ.

**FIG. 12.**
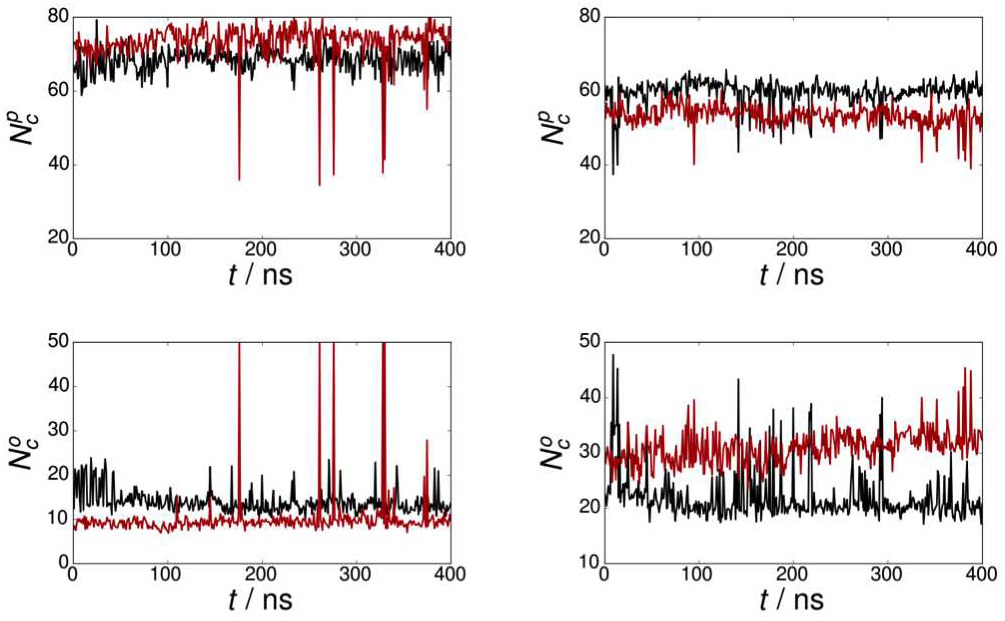
Number of contacts between hydrophobic residues (top) and hydrophobic residues and octane (bottom) for LSZ (left) and *α*La (right). Black and red lines denote Rotx and Rotz simulations.

## VII. CONCLUSIONS

Using atomistic molecular dynamics simulations the interfacial adsorption and conformation of two proteins, lysozyme and *α*-lactalbumin, was investigated. Due to their intrinsic amphiphilicity both proteins adsorb onto the interface. Running simulations with different starting protein orientation suggests that *α*La has a preferred orientation for adsorption to the interface, with adsorption occurring through regions near the membrane binding helices. By contrast LSZ adsorbs in a non-specific manner.

Due to their strong intramolecular interactions, including a number of disulfide bonds neither protein undergoes significant changes in structure during this initial adsorption. Using REST simulations we showed that both proteins exhibit compact and extended states. The extended states are expanded in the plane of the interface. For both proteins the compact states are more common than the extended states, in contrast to recent work investigating myoglobin derived peptides; this reflects the rigidity of these proteins.

Examining the different interfacial conformations reveals some differences between LSZ and *α*La. While compact states are more common for both proteins, for LSZ the interfacial conformations are dominated by a single conformation, with a variety of different extended states are also found. Using a cluster analysis 18 distinct interfacial conformations were found for LSZ and estimated free energy difference between the compact and extended states to be ≳ 3 kcal mol^−1^. *α*La has fewer conformations but these are populated more evenly. Again it is most commonly found in compact but has two highly populated compact states. The extended states are more highly populated than for LSZ with the free energy difference between the compact and extended states being smaller than for LSZ (~ 1.9 kcal mol^−1^).

As would be expected the extended states are stabilised by the decrease in interfacial free energy, due to the increase in occupied interfacial area, and the partitioning of hydrophobic residues into the oil phase. The compact states are stabilised by the protein intramolecular interactions. For *α*La the extended states are additionally stabilised by contacts between hydrophobic residues and octane molecules, leading to a decrease in the protein-octane interaction energy. LSZ in contrast shows no significant difference in protein-octane interaction energy between the compact and extended states. Indeed examining the joint probability distribution *P*(*R*_*g*_, *E*_*po*_) showed that the lowest protein-octane interaction energy is found for a cluster of compact states.

This insight into the conformations adopted by proteins at interfaces can give new insight into protein interfacial adsorption and aggregation. The contrast between the behaviour of these proteins and myoglobin-derived peptides studied in recent work demonstrates the key role that protein intramolecular interactions play in determining interfacial conformations. While LSZ and *α*La are largely found in compact states, the weaker intermolecular interactions in the myoglobin peptides leads to the formation of extended states^33^. Weakening the interactions in LSZ and *α*La, for example through reducing some the disulfide bonds, may be used to promote the formation of extended states.

One noticeable difference in behaviour between LSZ and *α*La is the interaction between the protein and oil molecules. For *α*La the protein-oil interaction is more favourable in the extended state while no significant difference was found for LSZ. This suggests that the effect of the oil phase on the interfacial conformation is dependent on the protein. The destabilisation of the compact state by protein-oil interactions is consistent with recent work on the protein biosurfactant Rsn-2, where the clamshell opening was found at the cyclohexane-water interface but not at the air-water interface due to penetration of oil molecules into the hydrophobic protein core^22^.

While SRCD studies showed that the proportion of *α*-helix for *α*La increased at the water-alkane interface this was not seen in our simulations. This could be a timescale issue, as the continuous simulation time was 900 ns, which may still not be long enough to observe secondary structure change for these larger proteins. The SRCD experiments were performed at a water-hexadecane interface while we used a water-octane interface in this work; however, as an alkane was used in both cases this is unlikely to have a significant effect. Alternatively interactions between proteins at the interface could also cause changes in the protein secondary structure. The protein interfacial density in this work is about a factor of 5 lower than the lowest density examined in experiment. This could also explain the differences in secondary structure for LSZ; while we see a decrease in the *α*-helix proportion we do not see an increase in *β*-sheet proportion. Aggregation of LSZ at the interface could lead to the formation of intermolecular *β*-sheets that we were unable to observe in this work.

## ACKNOWLEDGEMENTS

I wish to acknowledge the SFI/HEA Irish Centre for High-End Computing (ICHEC) for the provision of computational facilities and support. The modified version of PLUMED for performing REST simulations was provided by Giovanni Bussi (SISSA).

## APPENDICES

### Appendix A Convergence of REST simulations

The convergence of the REST simulations was monitored through the variation in the REST scaling factor (*β*_*i*_) for each of the replicas. Shown in Figure 13 plots of At *t* = 0 the *j*th replica starts at *β* = *β*_*j*_. As can be seen the majority of the replicas explore the different values of *β*_*i*_. Most notably the replica in the *β*_0_ state, which is the only physically relevant replica, changes regularly across the simulation.

**FIG. 13.**
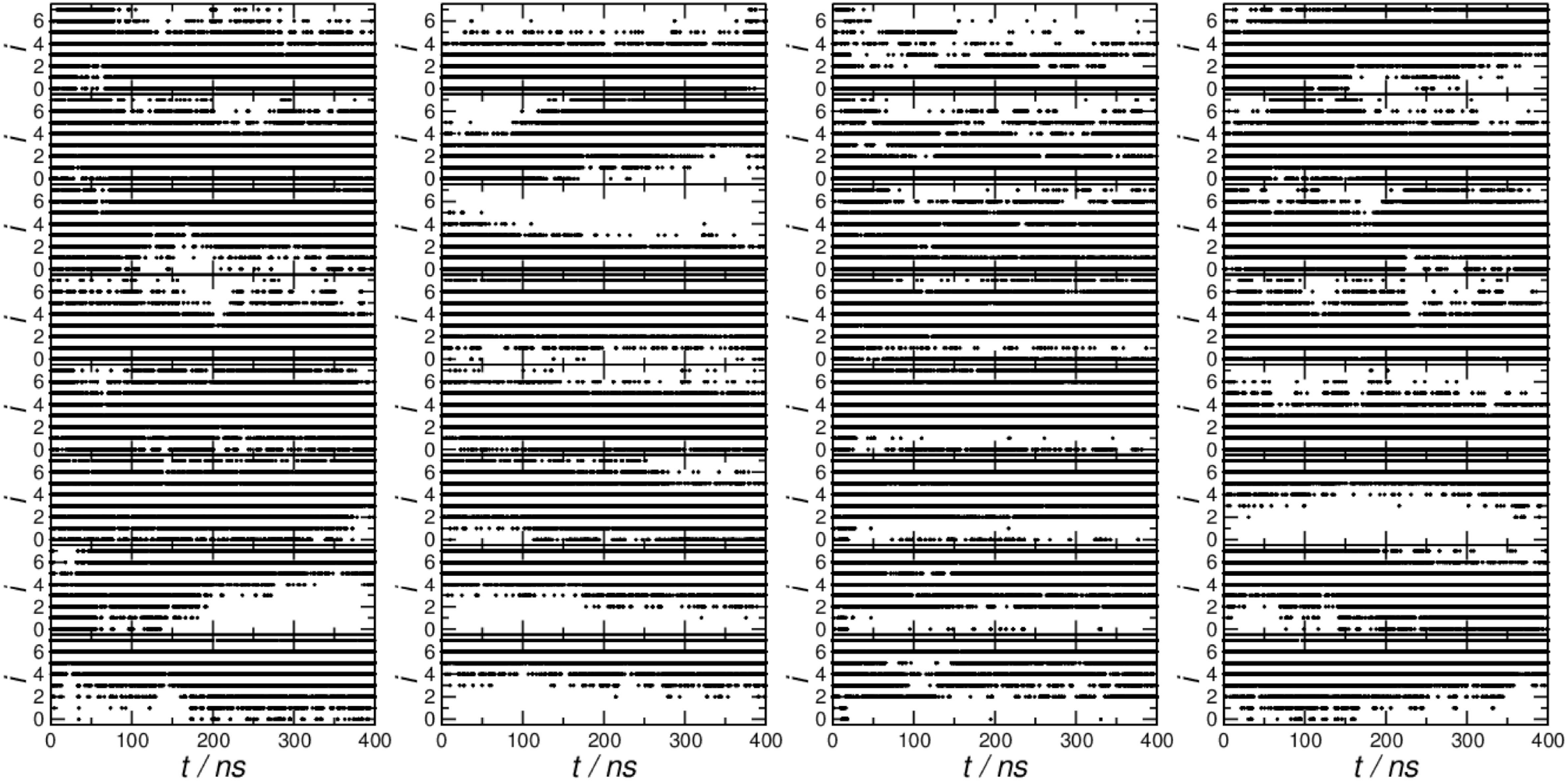
Plot of scaling factor (*i* denotes *β*_*i*_) against time for (left to right) LSZ Rotx, LSZ Rotz, *α*La Rotx, and *α*La Rotz, for different simulation replicas. From top to bottom graphs show replicas with *i*=0, 1, 2, 3, 4, 5, 6, and 7 at *t*=0 ns.

**FIG. 14.**
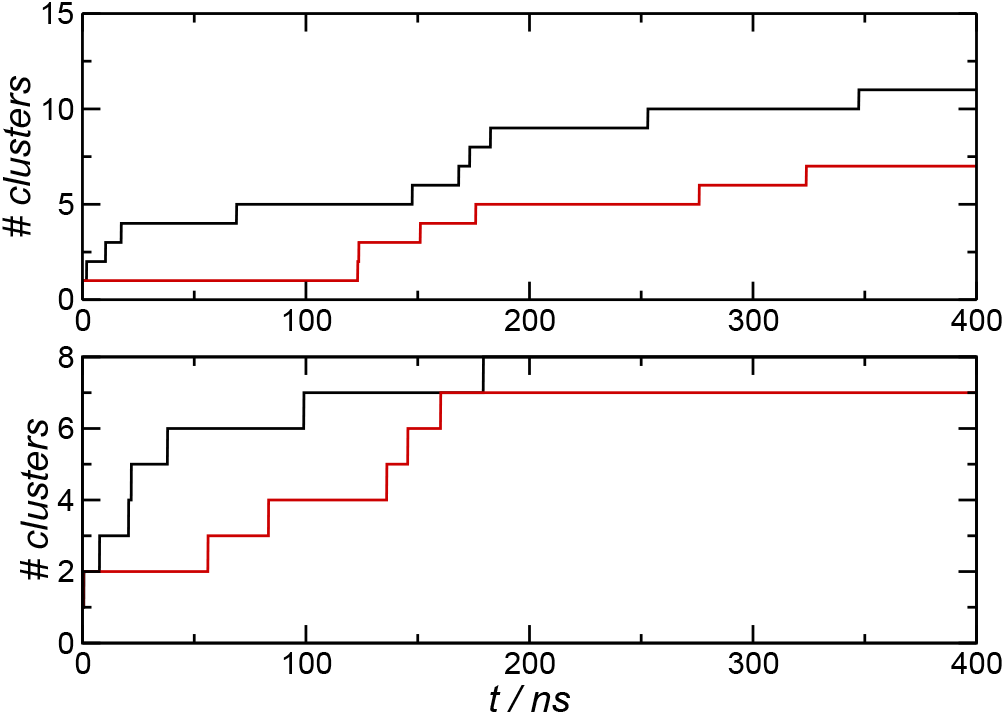
Variation in number of distinct clusters with time across the REST simulations for LSZ (top) and *α*La (bottom). Rotx and Rotz simulations denoted by black and red lines respectively.

Additionally the number of distinct conformations found in the REST simulations as a function of time was determined. As this was performed over the entire length of the simulations and separately for the Rotx and Rotz simulations for both proteins the number clusters are different than in Sec. V. After a relatively rapid increase at the beginning of the simulations this is increases only slowly (for LSZ) or remains constant (for *α*La) across the last 100 ns.

### Appendix B Radius of gyration probability maps

Shown in Figure 15 are two-dimensional probability maps 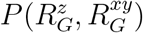 for LSZ and *α*La. Most of the conformations for LSZ fall into a single compact state and a scattered set of extended states (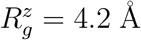 and 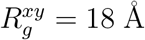). There is also a small cluster of conformations (around 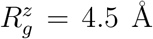 and 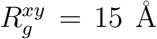) that may be considered intermediate between the compact and extended states. *α*La has two compact states and single set of extended states, clustered around 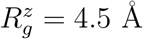 and 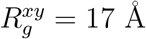.

**FIG. 15.**
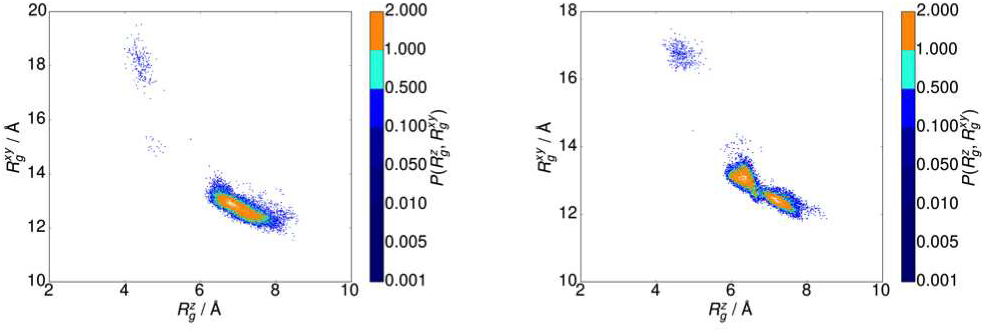
Probability maps for LSZ (left) and *α*La (right) from REST simulation (T=298 K).

## Reference

1 E. Dickinson, Soft Matter 2, 642 (2006).

2 N. R. Stanley-Wall and C. E. MacPhee, Phys. Biol. 12, 063001 (2015).

3 A. Schulz, B. M. Liebeck, D. John, A. Heiss, T. Subkowski, and A. Böker, J. Mater. Chem. 21, 9731 (2011).

4 A. Cooper and M. W. Kennedy, Biophys. Chem. 151, 96 (2010).

5 M. Schor, J. L. Reid, C. E. MacPhee, and N. R. Stanley-Wall, Trends Biochem. Sci. 41, 610 (2016).

6 M. B. Linder, Curr. Opin. Colloid Interface Sci. 14, 356 (2009).

7 A. Cooper, M. W. Kennedy, R. I. Fleming, E. H. Wilson, H. Videler, D. L. Wokosin, T.-J. Su, R. J. Green, and J. R. Lu, Biophys. J. 88, 2114 (2005).

8 S. J. Vance, R. E. Mcdonald, A. Cooper, B. O. Smith, and M. W. Kennedy, J. R. Soc. Interface 10, 20130453 (2013).

9 P. Reis, K. Holmberg, H. Watzke, M. E. Leser, and R. Miller, Adv. Colloid Interface Sci. 147–148, 237 (2008).

10 J. A. Zasadzinski, J. Ding, H. E. Warriner, F. Bringezu, and A. J. Waring, Curr. Opin. Colloid Interface Sci. 6, 506 (2001).

11 Y. F. Yano, J. Phys. Condens. Matter 24, 503101 (2012).

12 J. L. Zhai, L. Day, M.-I. Aguilar, and T. J. Wooster, Curr. Opin. Colloid Interface Sci. 18, 257 (2013).

13 C. D. Mackenzie, B. O. Smith, A. Meister, A. Blume, X. Zhao, J. R. Lu, M. W. Kennedy, and A. Cooper, Biophys. J. 96, 4984 (2009).

14 D. Gidalevitz, Z. Huang, and S. A. Rice, Proc. Natl. Acad. Sci. U. S. A. 96, 2608 (1999).

15 X. Zhao, F. Pan, and J. R. Lu, J. R. Soc. Interface 6, S659 (2009).

16 I. M. Tucker, J. T. Petkov, J. Penfold, R. K. Thomas, A. R. Cox, and N. Hedges, Langmuir 31, 10008 (2015).

17 F. A. Husband, M. J. Garrood, A. R. Mackie, G. R. Burnett, and P. J. Wilde, J. Agric. Food Chem. 49, 859 (2001).

18 A. J. Miles and B. A. Wallace, Chem. Soc. Rev. 35, 39 (2006).

19 S. Jordens, P. A. Rühs, C. Sieber, L. Isa, P. Fischer, and R. Mezzenga, Langmuir 30, 10090 (2014).

20 D. Zare, K. M. McGrath, and J. R. Allison, Biomacromolecules 16, 1855 (2015).

21 D. Zare, J. R. Allison, and K. M. McGrath, Biomacromolecules 17, 1572 (2016).

22 G. B. Brandani, S. J. Vance, M. Schor, A. Cooper, M. W. Kennedy, B. O. Smith, C. E. MacPhee, and D. L. Cheung, Phys. Chem. Chem. Phys. 19, 8584 (2017).

23 A. Schulz, M. Fioroni, M. B. Linder, A. Nessel, M. Bocola, T. Subkowski, U. Schwaneberg, A. Böker, and F. Rodríguez-Ropero, Soft Matter 8, 11343 (2012).

24 A. Jain, M. Jochum, and C. Peter, Langmuir 30, 15486 (2014).

25 D. L. Cheung, Langmuir 28, 8730 (2012).

26 K. M. Bromley, R. J. Morris, L. Hobley, G. Brandani, R. M. C. Gillespie, M. McCluskey, U. Zachariae, D. Marenduzzo, N. R. Stanley-Wall, and C. E. MacPhee, Proc. Natl. Acad. Sci. 112, 5419 (2015).

27 C. A. Miller, N. L. Abbott, and J. J. de Pablo, Langmuir 25, 2811 (2009).

28 O. Engin, A. Villa, M. Sayar, and B. Hess, J. Phys. Chem. B 114, 11093 (2010).

29 C. Dalgicdir, C. Globisch, C. Peter, and M. Sayar, PLOS Comput. Biol. 11, e1004328 (2015).

30 D. J. Earl and M. W. Deem, Phys. Chem. Chem. Phys. 7, 3910 (2005).

31 A. Laio and F. L. Gervasio, Reports Prog. Phys. 71, 126601 (2008).

32 M. Arooj, N. S. Gandhi, C. A. Kreck, D. W. Arrigan, and R. L. Mancera, J. Phys. Chem. B 120, 3100 (2016).

33 D. L. Cheung, Langmuir 32, 4405 (2016).

34 J. Zhai, A. J. Miles, L. K. Pattenden, T.-H. Lee, M. A. Augustin, B. A. Wallace, M.-I. Aguilar, and T. J. Wooster, Biomacromolecules 11, 2136 (2010).

35 J. Zhai, S. V. Hoffmann, L. Day, T.-H. Lee, M. A. Augustin, M.-I. Aguilar, and T. J. Wooster, Langmuir 28, 2357 (2012).

36 L. Day, J. Zhai, M. Xu, N. C. Jones, S. V. Hoffmann, and T. J. Wooster, Food Hydrocoll. 34, 78 (2014).

37 E. Gasteiger, C. Hoogland, A. Gattiker, S. Duvaud, M. Wilkins, R. Appel, and B. A., in Proteomics Protoc. Handb. (2005), pp. 571–601.

38 H. Schwalbe, S. B. Grimshaw, A. Spencer, M. Buck, J. Boyd, C. M. Dobson, C. Redfield, and L. J. Smith, Protein Sci. 10, 677 (2001).

39 A. C. Pike, K. Brew, and K. R. Acharya, Structure 4, 691 (1996).

40 W. Humphrey, A. Dalke, and K. Schulten, J. Mol. Graph. 14, 33 (1996).

41 D. van der Spoel, E. Lindahl, B. Hess, G. Groenhof, A. E. Mark, and H. J. C. Berendsen, J. Comput. Chem. 26, 1701 (2005).

42 B. Hess, C. Kutzner, D. van der Spoel, and E. Lindahl, J. Chem. Theory Comput. 4, 435 (2008).

43 G. A. Tribello, M. Bonomi, D. Branduardi, C. Camilloni, and G. Bussi, Comput. Phys. Commun. 185, 604 (2014).

44 G. Bussi, Mol. Phys. 112, 379 (2013).

45 N. Schmid, A. P. Eichenberger, A. Choutko, S. Riniker, M. Winger, A. E. Mark, and W. F. van Gunsteren, Eur. Biophys. J. 40, 843 (2011).

46 H. J. C. Berendsen, J. R. Grigera, and T. P. Straatsma, J. Phys. Chem. 91, 6269 (1987).

47 C. Oostenbrink, A. Villa, A. E. Mark, and W. F. van Gunsteren, J. Comput. Chem. 25, 1656 (2004).

48 S. R. Euston, P. Hughes, M. A. Naser, and R. E. Westacott, Biomacromolecules 9, 1443 (2008).

49 S. R. Euston, Food Hydrocoll. 42, 66 (2013).

50 U. Essmann, L. Perera, M. L. Berkowitz, T. Darden, H. Lee, and L. G. Pedersen, J. Chem. Phys. 103, 8577 (1995).

51 G. Bussi, D. Donadio, and M. Parrinello, J. Chem. Phys. 126, 014101 (2007).

52 B. Hess, H. Bekker, H. J. C. Berendsen, and J. G. E. M. Fraaije, J. Comput. Chem. 18, 1463 (1997).

53 P. Liu, B. Kim, R. A. Friesner, and B. J. Berne, Proc. Natl. Acad. Sci. U. S. A. 102, 13749 (2005).

54 Y. Sugita and Y. Okamoto, Chem. Phys. Lett. 314, 141 (1999).

55 N. Michaud-Agrawal, E. J. Denning, T. B. Woolf, and O. Beckstein, J. Comput. Chem. 32, 2319 (2011).

56 D. Frishman and P. Argos, Proteins Struct. Funct. Genet. 23, 566 (1995).

57 X. Daura, K. Gademann, B. Jaun, D. Seebach, W. F. van Gunsteren, and A. E. Mark, 38, 236 (1999).

58 R. L. C. Vink, J. Horbach, and K. Binder, J. Chem. Phys. 122, 134905 (2005).

59 A. Radzicka and R. Wolfenden, Biochemistry 27, 1664 (1988).

60 Ø. Halskau, N. Å. Frøystein, A. Muga, and A. Martínez, J. Mol. Biol. 321, 99 (2002).

61 M. Schiffer and A. B. Edmundson, 7, 121 (1967).

62 W. C. Wimley and S. H. White, Nat. Struct. Mol. Biol. 3, 842 (1996).

63 S. Zeppieri, J. Rodríguez, and A. L. López de Ramos, J. Chem. Eng. Data 46, 1086 (2001).

